# Natural Epiallelic Variation is Associated with Quantitative Resistance to the Pathogen *Plasmodiophora Brassicae*

**DOI:** 10.1101/776989

**Authors:** Benjamin Liégard, Antoine Gravot, Leandro Quadrana, Yoann Aigu, Juliette Bénéjam, Christine Lariagon, Jocelyne Lemoine, Vincent Colot, Maria J. Manzanares-Dauleux, Mélanie Jubault

## Abstract

Clubroot caused by the protist *Plasmodiophora brassicae* is a major disease affecting cultivated *Brassicaceae*. Here, we uncover the existence of a natural epigenetic variation that is associated with partial resistance to clubroot in *Arabidopsis,* by using QTL fine mapping followed by extensive DNA sequence and methylation analyses. We show that at QTL *Pb-At5.2*, DNA methylation variation is extensive across accessions and strictly correlates with expression variation of the two neighboring genes At5g47260 and At5g47280, which encode NLR-immune receptors. Moreover, these natural variants are stably inherited and are not consistently associated with any nucleotide variation. These findings suggest a direct role for epigenetic variation in quantitative resistance of plants to pathogen attacks.

---

Intraspecific diversity in plant immune interactions is associated with the high level of sequence variations at hundreds of NLR, one of the largest and most rapidly evolving plant gene families (Stahl et al., 1999; Meyers et al., 2003; Clark et al., 2004; Innes et al., 2008; Yue et al., 2012; Shao et al., 2016). Each NLR protein can be involved in the direct or indirect recognition of a small range of effector proteins secreted by specific strains of plant pathogens, potentially triggering the induction of strong plant defense responses that can rapidly stops pathogen invasion (Maekawa et al., 2011; Jones et al., 2016). Thus, with only a few exceptions (including non-NLR driven resistances (Thomas, 1998; Xiao et al., 2001; Larkan et al., 2013) and NLR-driven broad-spectrum resistances (Ernst et al., 2002; Qu et al., 2006)), the catalogue of NLR encoding *R*-genes expressed in a given plant genotype shapes the range of isolate-specific full resistances (incompatible interactions).

In contrast to *R*-gene driven resistance, quantitative resistance is polygenic, i.e. it involves allelic variation at several Quantitative Trait Loci (QTL), which collectively contribute to post-invasive partial resistance in compatible plant-pathogen interactions. An emerging list of cloned resistance QTL supports the premise that Quantitative Resistance Genes (QRG) are functionally more diverse than *R* genes (Pilet-Nayel et al., 2017; Nelson et al., 2017).This short list however still includes genes encoding NLR (Hayashi et al., 2010; Fukuoka et al., 2014; Xu et al., 2014; Debieu et al., 2016) and other receptors (Diener and Ausubel, 2005; Hurni et al., 2015) or co-receptors (Huard-Chauveau et al., 2013). Thus, variation in NLR genes (or other non-self-recognition loci) appears to contribute to the variations in basal resistance levels in compatible interactions.

The triggering of effective resistance requires that cellular levels of NLR proteins reach minimum thresholds. However, elevated expression levels of NLR can also lead to autoimmunity drawbacks, including spontaneous HR and retarded plant growth (Li et al., 2015; Lai and Eulgem, 2018). Their abundance is thus tightly controlled by multiple mechanisms, at the transcriptional, post-transcriptional (Zhang and Gassmann, 2007) (*i.e.* alternative splicing) and post-translational levels (*i.e.* Ubiquitin-dependent proteolytic regulation). NLR regulation also involves a wealth of epigenetic-related cellular processes, including redundant networks of small RNA (Shivaprasad et al., 2012; Fei et al., 2013; Deng et al., 2018), histone modifications (Palma et al., 2010; Xia et al., 2013; Zou et al., 2014), histone-mark dependent-alternative splicing (Tsuchiya and Eulgem, 2013), regulation of chromatin structure and DNA methylation (Li et al., 2010). There is increasing evidence that epigenetic processes can play roles in the transitory imprinting of some plant biotic stress responses, at least for a few generations (Molinier et al., 2006; Slaughter et al., 2011; Luna et al., 2012). It is however not yet clear to which extent the stable transgenerational inheritance of epigenetically regulated gene expression contributes to the natural intraspecific diversity of plant-pathogen interactions.

The few available examples of transgenerational epigenetically controlled traits are mostly found in plant species, where the association between natural or induced differentially methylated regions (DMR) and phenotypic traits were shown stably or (most often) metastably inherited across the generations (Quadrana and Colot, 2016; Furci et al., 2019; Liégard et al., 2019). Such regions, designated as epialleles, can affect agronomically relevant traits: compatibility, accumulation of vitamin E and fruit ripening in tomato, disease resistance, sex determination in melon, and fruit productivity in oil palm.

In plants, DNA methylation can occur at cytosines in the three sequence contexts, CG, CHG and CHH (Henderson and Jacobsen, 2007) (where H could be A, C or T) and its impact varies depending on the targeted genomic features (i.e. transposable elements, gene promoters or gene bodies). DNA methylation patterns result from the dynamic combination of *de novo* methylation, maintenance methylation and demethylation. The *de novo* DNA methylation is catalyzed by the canonical and non-canonical RNA-directed DNA methylation (RdDM) pathways, which are both guided by siRNA (Zhang et al., 2018). Maintenance of DNA methylation mainly relies on RNA-independent pathways and require the activity of DDM1, MET1 and VIM proteins at CG sites, and of DDM1, KYP, CMT2/3 and histone mark HK9me2 at CHG and CHH sites (Law and Jacobsen, 2010; Matzke and Mosher, 2014)). Previous studies highlighted that natural DMRs were overrepresented over genes belonging to the *NLR* disease resistance gene family (Kawakatsu et al., 2016). However, it remains unclear if natural epigenetic variation at *NLR* genes can shape plant pathogen interactions.

Here, we report the identification of naturally occurring stable epigenetic variation underlying a QTL involved in partial resistance to clubroot in *Arabidopsis*. Clubroot is a root gall disease caused by the telluric biotrophic pathogen *Plasmodiophora brassicae* (Rhizaria), affecting all *Brassicaceae* crops. The infection process involves a primary infection in root hairs for only a few days. Then secondary plasmodia develop in root cortical cells, causing hyperplasia and hypertrophy that ultimately impair plant water and nutrient uptake. The reference accession Col-0 and Bur-0 are fully susceptible and partially resistant to *P. brassicae* isolate eH, respectively (Alix et al., 2007; Jubault et al., 2008b) (**Supplementary Figure S1**). Four main QTL determine this difference, which act additively^43^. Here we identify by fine mapping of the largest effect resistance QTL *Pb-At5.2* a strong association between partial resistance and the expression level of the two *NLR* genes *At5g47260* and *At5g47260,* linked to the DNA methylation status of the small region including those two genes and a neighboring transposable element (TE) sequence. Furthermore, we show that epiallelic variation at this locus is frequent among natural *Arabidopsis* accessions and that the highly methylated state is negatively correlated with the expression of the two *NLR*-genes as well as with quantitative resistance to *P. brassicae*. In contrast, there is no correlation between the trait and any specific nucleotide variant across the entire QTL interval. We further show that the RNA independent pathway involving DDM1, MET1, VIM and CMT2/3 maintains the hypermethylated epiallele in Col-0. Overall, our findings demonstrate that the quantitative resistance to a major root disease affecting *Brassicaceae* is associated, in *Arabidopsis*, with the stable inheritance of a natural epigenetic variation involved in the control of the constitutive expression of a pair of NLR-genes.

## Results

### Fine mapping of the Pb-At5.2 locus responsible for clubroot resistance

Using a population of F7 recombinant inbred lines (RILs) between the partially resistant accession Bur-0 and the susceptible Col-0, we previously mapped a QTL (*Pb-At5.2),* located on chromosome 5 between 67.5 and 71.8 cM, explaining a significant fraction (R^2^=20 %) of the resistance (**Fig. 1a**) (Alix et al., 2007; Jubault et al., 2008b). This interval contained 157 annotated sequences between At5g46260 and At5g47690. The effect and confidence interval of this QTL was also previously confirmed in Heterogeneous Inbred Family (HIF) lines 10499 and 13499 (Lemarié et al., 2015), both derived from the RIL 499 which harbored residual heterozygosity in the *Pb-At5.2* region (**Fig. 1b-c**, **Supplementary Text S1)**. The initial aim of the present work was to fine map *Pb-At5.2*, starting with reciprocal crosses between HIF lines 10499 and 13499. Clubroot symptoms in individuals of the F1 progeny were equally severe to those in the susceptible parental line HIF 13499, suggesting that the resistance allele Bur-0 was recessive (**Supplementary Figure S2**). The boundaries of the *Pb-At5.2* resistance locus was further refined through several rounds of genotyping and clubroot phenotyping (F3 to F5 generations downstream 10499/13499 crosses), details in **Fig. 1d-f**, **Supplementary Figure S3, Supplementary Text S1, Supplementary Data 1**). This enabled us to narrow down the confidence interval to 26 kb between the markers CLG4 (19,182,401 bp, in the promoter region of At5g47240), and the marker K64 (19,208,823 bp, in At5g47330). This region contained eight annotated open reading frames (ORFs), including the three NLR-encoding genes At5g47250/At5g47260/At5g47280, six annotated TE sequences and one lncRNA gene (**Fig. 1f**). The two F5 homozygous progeny lines 1381-2 and 2313-15, harboring the closest recombination events from both sides of the 26 kb interval, (see **Fig. 1e**) also showed partial resistance to a series of additional *P. brassicae* isolates (from pathotypes 1, 4 and 7 following the classification of Some et al. (1996). This highlighted the broad spectrum of the resistance conferred by the Bur-0 allele of *Pb-At5.2* (**Supplementary Figure S4**).

**Fig. 1.**
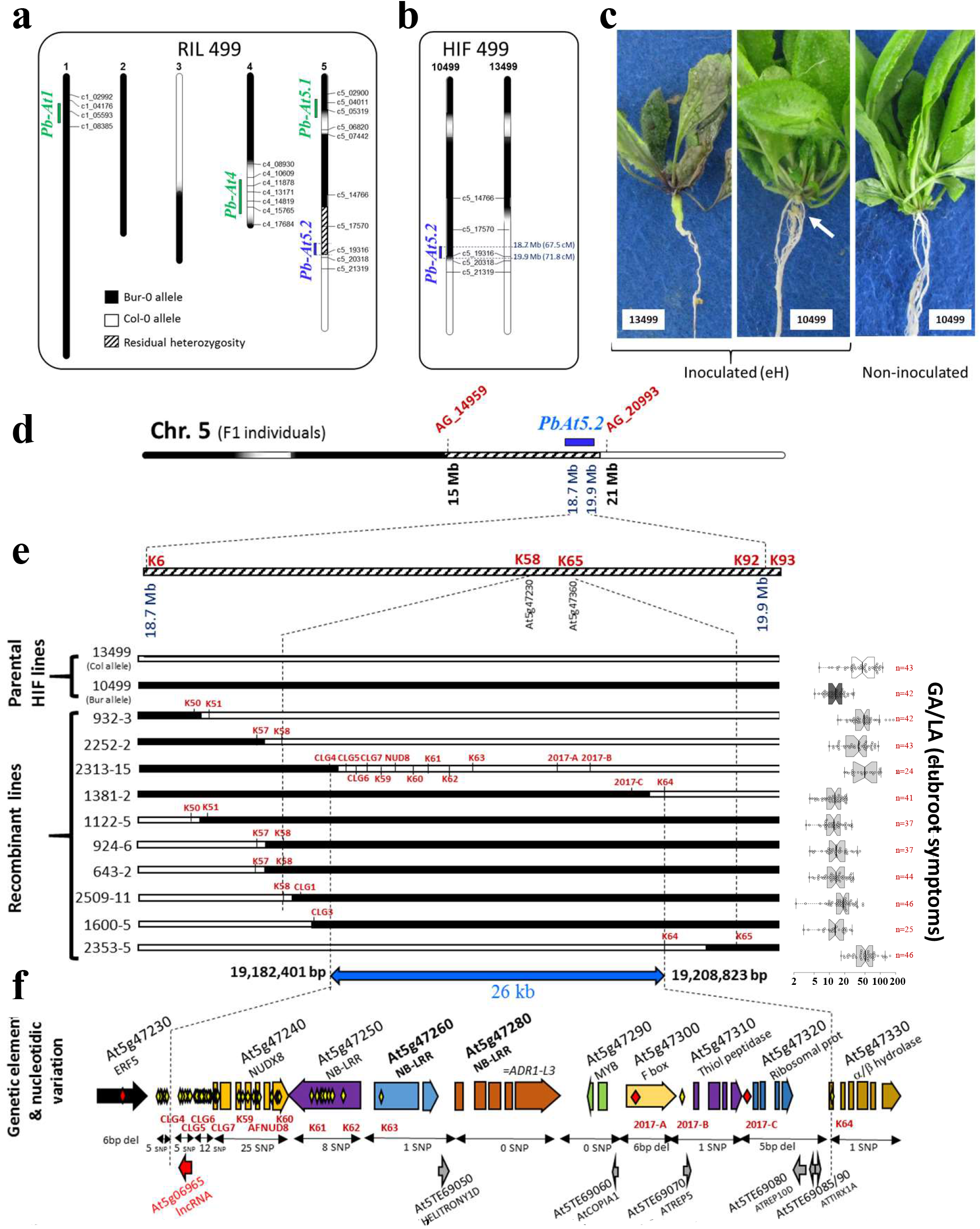
Fine mapping of *Pb-At5.2*. **a** Genetic map and residual heterozygosity in the Recombinant Inbred Line (RIL) 499 derived from Bur-0 and Col-0 and position of clubroot resistance QTL (from Jubault et al. (2008b)). Black: Bur-0 allele, White: Col-0 allele, Hatched: Heterozygous (Col-0/Bur-0). **b** Allele configuration at *Pb-At5.2* in the two derived HIF lines 10499 and 13499. **c** Photos showing *Pb-At5.2* conferred partial resistance to the eH isolate conferred by *Pb-At5.2* in the RIL499 genetic background. Observations were made at 21 days post-inoculation. The white arrow indicates the presence of a limited number of galls in inoculated 10499. **d** First round of fine mapping: allelic structure in the F1 lines derived from reciprocal crosses between 10499 and 13499. 554 F3 lines with recombination in the confidence interval were screened using 10 SNP markers between AG_14959 and AG_20993. High density genotyping (94 SNP from K1 to K94), in a series of 88 recombinant F3 lines, and clubroot phenotyping in their bulked segregating F4 progenies, led to a new interval between markers K58 (At5g47230) and K65 (At5g47360). **e** Second round of fine mapping: Recombination positions in homozygous individuals obtained from selected recombinant lines. For each line, the GA/LA index (disease symptoms) is indicated on the right panel. Center lines show the medians; box limits indicate the 25th and 75th percentiles as determined by R software; whiskers extend 1.5 times the interquartile range from the 25th and 75th percentiles, outliers are represented by dots; data points are plotted as open circles. Number of individual plants analysed for each genotype is indicated (n). The notches are defined as +/−1.58*IQR/sqrt(n) and represent the 95% confidence interval for each median. Genetic markers are indicated for each recombination position. Markers between CLG5 and 2017-C were used in every line, but only shown for 2313-15. **f** New 26 kb interval of *Pb-At5.2* between markers CL4 (excluded) and K64 (excluded), containing eight annotated ORF, six transposons and one lncRNA. Yellow and red diamonds indicate SNP and nucleotide deletions, respectively.

### Possible causal role for a stably inherited expression/methylation polymorphism affecting two NLR genes

RNA-seq analysis was carried out on the Bur-0 and Col-0 accessions and the recombinant HIF lines 10499 and 13499. Pathogen-induced gene patterns markedly differed in genotypes harboring alleles *Pb-At5.2*_*BUR*_ or *Pb-At5.2*_*COL*_ (**Supplementary Figure S5, Supplementary Data 2**). Those regulations were consistent with our previously published studies *i.e.* a role for camalexin biosynthesis and SA-mediated responses in *Pb-At5.2*_*BUR*_-mediated resistance; a role for JA-driven induction of *ARGAH2* in *Pb-At5.2*_*COL*_-mediated basal resistance (details in **Supplementary Text 2**). We then focused on the eight ORFs in *Pb-At5.2*. In Col-0, sequenced reads were only found for four of the genes: At5g47240/At5g47250/At5g47310/At5g47320. At5g47310 and At5g47320 only had one SNP and one indel in their promoter regions in Bur-0, respectively. The possible causal role of these variations was discarded, as both genes were expressed similarly from the Bur-0 and Col-0 alleles (**Fig. 2a**). Most of the other SNPs in the 26kb region were associated with the NUDX8 encoding gene At5g47240 and to a smaller extent with the adjacent NLR-gene At5g47250. However, clubroot symptoms in the homozygous T-DNA mutant lines SALK_092325C (T-DNA in the At5g47240 gene) and WiscDsLoxHs110_09B (T-DNA in the At5g47250 gene) were the same as in Col-0 (**Supplementary Figure S6**). At5g47260 and At5g47280, both encoding proteins belonging to the family of non-TIR-NLR immune receptors, displayed one single non-synonymous SNP, and no SNP, respectively. However, RNA-seq analysis indicated that these two genes were constitutively expressed in the roots of Bur-0, 10499 and 1381-2 (with the Bur-0 allele, **Fig. 2a** and **Supplementary Figure S4**), but their expression was almost undetectable in Col-0, 13499 and 2313-15 (with the Col-0 allele).

**Fig. 2.**
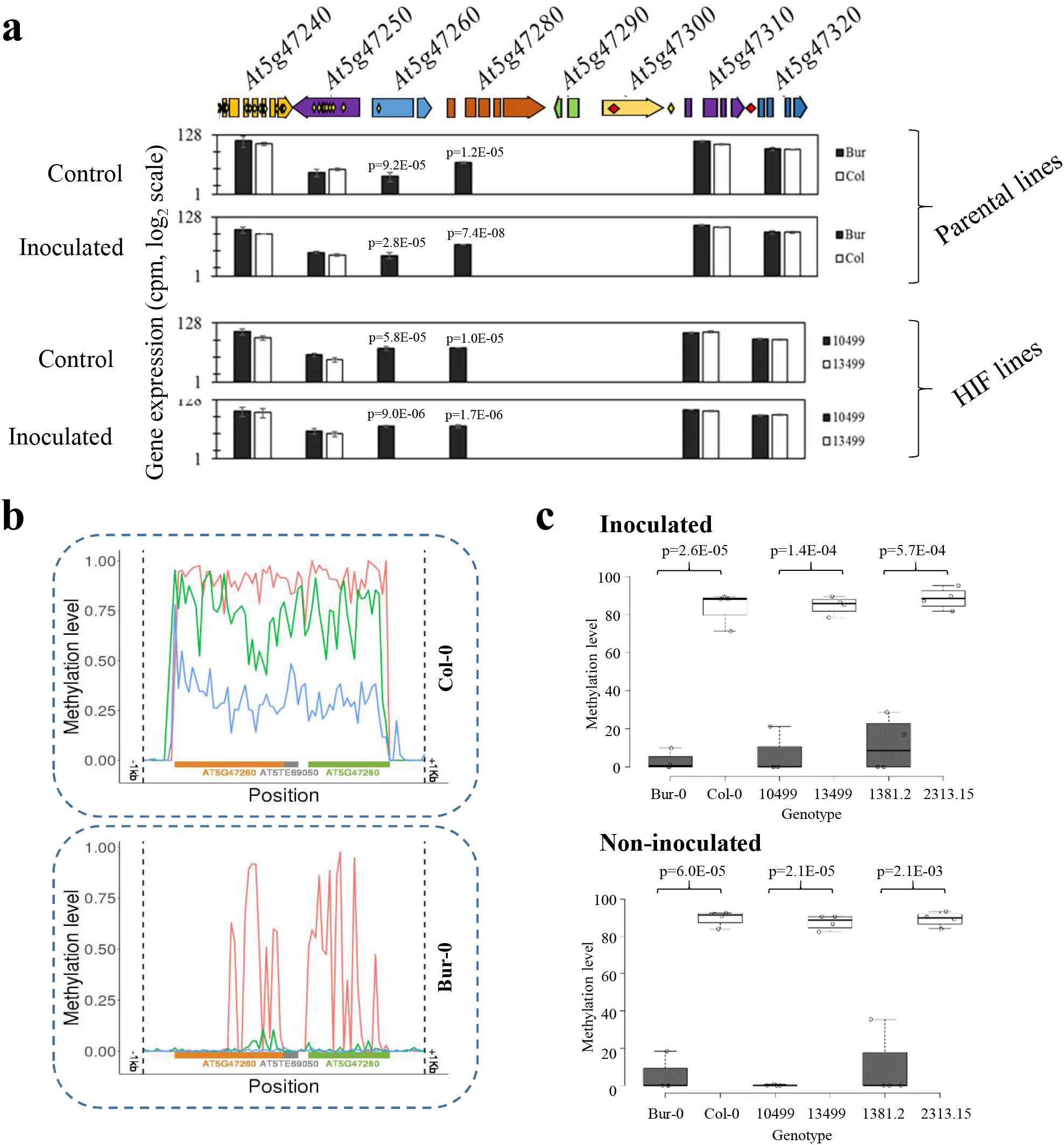
Identification of two candidate NLR-encoding genes with contrasted expression and methylation profiles at QTL *Pb-At5.2*. **a** Sequence variations and expression levels of genes in the *Pb-At5.2* region. Yellow and red diamonds indicate SNP and INDEL variations, respectively. Gene expression values are from RNAseq analyses conducted in inoculated and control conditions at 14 dpi (log2 normalized cpm), with parental lines Col-0/Bur-0 and HIF lines 13499/10499 (the last two were derived from the RIL 499, and homozygous *Pb-At5.2_Col/Col_* and *Pb-At5.2*_Bur/Bur_, respectively) (**Supplementary Data 2**). FDR adjusted *p-values* are shown if less than 0.05. **b** Methylation profiles in the At5g47260 and At5g47280 region, in Col-0 and Bur-0 accessions, inferred from bisulfite data previously reported in Kawakatsu et al. (2016) Average methylation level was calculated within non-overlapping 100-bp windows starting 1 kb before the TSS site of At5g47260 and stopping 1 kb after the TSE site of At5g47280. In red: methylation in the CG context. In green: methylation in the CHG context. In blue: methylation in the CHH context. **c** Methylation profiles obtained by CHOP-qPCR on At5g47260 in inoculated and non-inoculated roots of Bur-0/Col-0, 10499/13499, and in the homozygous recombinant lines 2313-15 (*Pb-At5.2*_Col/Col_) and 1381-2 (*Pb-At5.2*_Bur/Bur_). Those two last genotypes harbor the narrowest recombination events from either side of *Pb-At5.2* (between markers CLG4 and K64, details in Fig. 1). Center lines show the medians; box limits indicate the 25th and 75th percentiles as determined by R software; whiskers extend 1.5 times the interquartile range from the 25th and 75th percentiles, outliers are represented by dots; data points are plotted as open circles. n = 4 bulks of 6 plants and *p-values* are shown (sided t-test).

To understand why these two NLR genes At5g47260 and At5g47280 were differentially expressed in Bur-0 and Col-0, even in the absence of any sequence variation within the putative promoter regions of these genes, we analyzed the DNA methylation level at the region in these two accessions using public methylome data (Kawakatsu et al., 2016). The genomic interval between 19,188,411 and 19,196,559, which includes, in Col-0 and Bur-0, the two genes At5g47260 and At5g47280 and the transposon At5TE69050 in-between, were hypermethylated and hypomethylated in Col-0 and Bur-0, respectively (**Fig. 2b**). This contrasting methylation state was experimentally confirmed using DNA extracted from infected or control roots of Col-0 and Bur-0 plants (**Fig. 2c**) and CHOP qPCR. These DNA methylation differences were also found between the progeny HIF lines 10499 and 13499 and in the pair of HIF-derived homozygous near-isogenic lines 1381-2/2313-15 (**Fig. 2c**), thus indicating that they are stably inherited independent of any DNA sequence polymorphism outside of the locus. Moreover, the ‘Col-like’ hypermethylation of At5g47260 and At5g47280 was systematically associated with a low expression of the two NLR genes and a lower level of partial resistance to *P. brassicae* infection. To further investigate the inheritance of this epiallelic variation and its penetrance on gene expression and clubroot resistance, we then investigated two series of 100 individual plants, corresponding to the progenies derived from selfing the heterozygous 2509 and 1381 lines (harbouring heterozygosity at the locus). The evaluation of plant disease for each individual plant in the two progenies indicated a 3:1 mendelian segregation of the partial resistance phenotype. Clubroot symptoms in individuals with only one Bur-0 resistance allele were the same as in individuals with the two susceptible Col-0 allele (**Fig. 3a**). In clubroot-inoculated roots of each of those individual plants from the 2509 progeny, the methylation state of the *Pb-At5.2* region was monitored by CHOP qPCR on At5g47260. In addition, the SNP allele status at *Pb-At5.2* was investigated for each individual plant (details of markers are given in **Supplementary Data 1**). Heterozygous Bur/Col individuals displayed intermediate parental methylation and expression values (**Fig. 3b-d**), thus providing a molecular explanation for the recessivity of the Bur-0 resistance allele. Altogether, these results suggested a link between partial resistance to *P. brassicae* and a stably inherited epiallelic variation at *Pb-At5.2*, which controls the expression of two NLR genes.

**Fig. 3.**
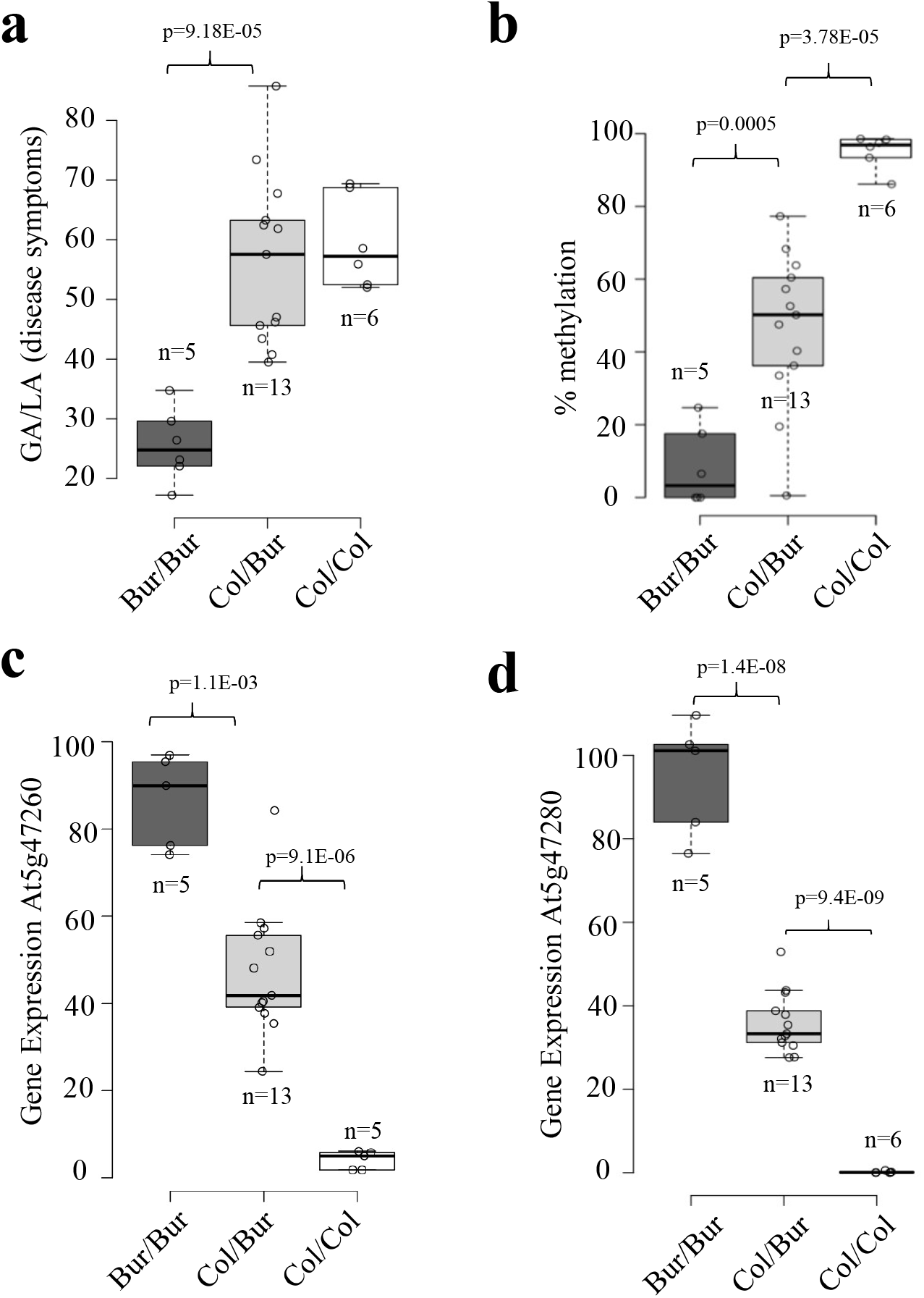
Intermediate methylation and transcription levels of candidate genes in heterozygous plants are associated with full clubroot susceptibility. Eighty-three individual plants of the segregating progeny from the recombinant line 2509 (heterozygous in Chr.5 regions between genetic markers K58 and K93) were sampled at 21 dpi. Leaves from each individual plant were used for genotyping (PCR marker CL_N8), which defined n pools of >3 plants of each zygosity profile: Bur/Bur (n=5), Col/Bur (n=13) and Col/Col (n=6) (black, grey and white boxes, respectively). Each plant pool was evaluated for **a** clubroot resistance (GA/LA), **b** % methylation at the locus, and **c-d** candidate gene expression (At5g47260 and At5g47280). Gene expression was quantified through RT-qPCR, data were normalized over mean-Cp from the pools Bur/Bur, following Pfaffl’s method with two reference genes (Pfaffl, 2001). Center lines show the medians; box limits indicate the 25th and 75th percentiles as determined by R software; whiskers extend 1.5 times the interquartile range from the 25th and 75th percentiles, outliers are represented by dots; data points are plotted as open circles.

### The Bur-like hypo-methylated epiallele is well represented among Arabidopsis accessions and contributes to reduce clubroot symptoms

To assess the relative contribution of DNA sequence and DNA methylation changes at *Pb-At5.2* to the variable resistance to clubroot we investigated the natural allelic and epiallelic diversity across *Arabidopsis* accessions. We took advantage of recently published Illumina short genome sequence reads obtained from 1135 Arabidopsis accessions (1001 Genomes Consortium, 2016) to document the species-wide molecular diversity of the *Pb-At5.2* genomic region. Based on quantitative horizontal and vertical coverage of short-reads aligned to the Col-0 reference genome sequences we identified two discrete groups of accessions. One of such groups, containing 401 accessions, was characterized by high vertical and horizontal coverage (>0.75) and included the reference accession Col-0 as well as the partially resistance Bur-0 (**Figure 4a**; detailed list of genotypes in **Supplementary Data 3, sheet 1**). Conversely, the remaining 734 accessions present diverse structural rearrangements, principally long deletions that translates in poor horizontal and vertical coverage compared to the reference Col-0 genome. Closer examination of coverage plots for the 401 accessions with Col-0/Bur-0-like revealed a uniform haplotype structure, which was present at high frequency at the species level (Minor Allele Frequency MAF ~0.37). Nonetheless, the haplotype frequency varied depending on geographic groups, ranging from 52.7 % in Spain to 17.7 % in Asia (**Supplementary Figure S7**). We then analyzed DNA methylation levels in 287 accessions belonging to the 401 accessions containing the Col-0/Bur-0-like *Pb-At5.2* and for which public bisulfite data is available (Kawakatsu et al., 2016). Based on this data, we could distinguish a group of 228 accessions showing hypomethylation of *Pb-At5.2*, including Bur-0, and another group of 59 accessions, which includes Col-0, that displays hypermethylation (**Fig. 4b**). The prevalence of accessions showing the Col-like (epi)haplotype varied considerably depending on geographic origin, ranging from 1.8 % in Spain to 16.8 % in Central Europe (**Supplementary Data 3, and Supplementary Figure S7**). Consistent with a causal role for DNA methylation in the transcriptional regulation At5g47260 and At5g47280, reanalysis of publicly available RNA-seq data revealed a pronounced negative correlation between methylation level and At5g47260 and At5g47280 gene expression (**Fig. 4c**). These results were further validated in infected roots of 20 natural accessions (**Fig. 4d**).

**Fig. 4.**
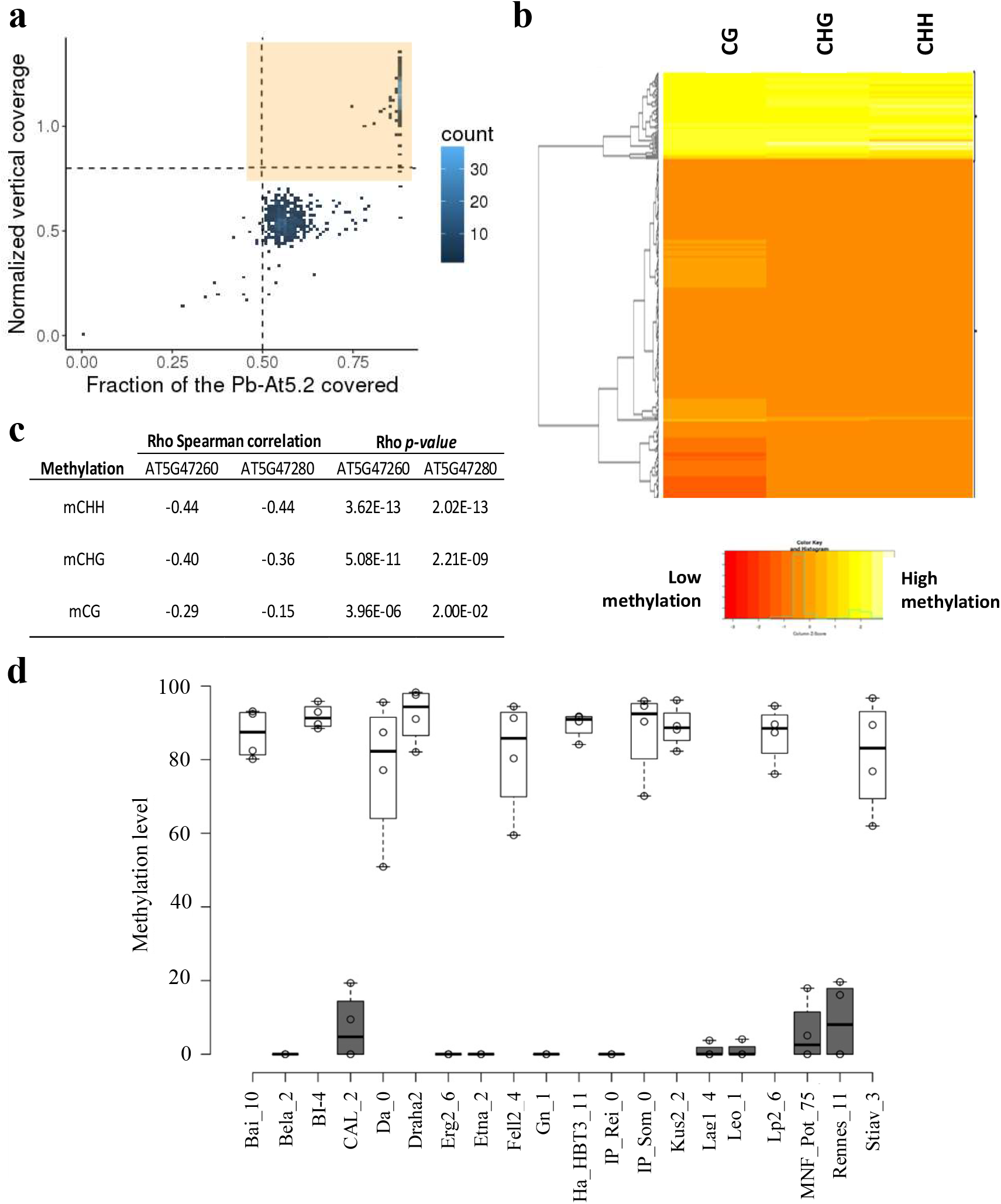
Natural epigenetic variation at *Pb-At5.2* affects the expression of At5g47260/Atg47280 among Arabidopsis accessions. **a** Screening for 1001 genome Arabidopsis accessions displaying a Col/Bur-like genomic structure at *Pb-At5.2* (chr5: 19185600 - 19200600). X axis: Horizontal coverage region covered by at least one read. Y axis: Vertical coverage in read percentage. The 401 accessions framed in the northeast intercardinal region delimited by dotted lines harbour a vertical read coverage >0.8 and an horizontal DNA-seq >0.5 (DNA-seq data from^46^). **b** Clustering of a series of accessions harbouring Col/Bur-like genomic structure at *Pb-At5.2*, by their level of methylation on At5g47260 and At5g47280. Bisulfite data were obtained from the 1001 genome data project **(Supplementary Data 3 sheet 2)**. Average methylation level was calculated 1 kb before the TSS site of At5g47260 and stopping 1 kb after the TSE site of At5g47280 for each context. **c** Spearman correlation between the methylation and gene expression of At5g47260 and At5g47280 in a subset of 253 *Arabidopsis* accessions for which expression data was available (RNAseq data from^41^). Correlation between gene expression and methylation level is given for all three DNA-methylation contexts in the interval from 1 kb before the TSS site of At5g47260, to 1 kb after the TSE site of At5g47280. **d** Confirmation of methylation profiles at At5g47260 in inoculated roots from 20 ecotypes. Methylation level was obtained using CHOP-qPCR. Black and white bars indicate genotypes with Bur-like and Col-like methylation patterns, respectively. Center lines show the medians; box limits indicate the 25th and 75th percentiles as determined by R software; whiskers extend 1.5 times the interquartile range from the 25th and 75th percentiles, outliers are represented by dots; data points are plotted as open circles. n = 4 bulks of 6 plants.

Both Col-like and Bur-like epialleles were significantly represented among natural accessions, thus offering interesting genetic material to determine the actual contribution of DNA sequence and DNA methylation in the control of clubroot partial resistance. A selection of 126 accessions, including 42 accessions with the Col-like epiallele and 85 accessions with the Bur-like epiallele, were assessed for their resistance to *P. brassicae* isolate eH. While no DNA sequence polymorphism within *Pb-At5.2* shows association with clubroot resistance (**Supplementary Data 4**), the low DNA methylation state of the At5g47260/At5g47280 locus was significantly associated with enhanced resistance levels (**Fig. 5**). Altogether, the results corroborate and extend the conclusions obtained by fine mapping of *Pb-At5.2* and provide strong evidence that natural epiallelic variation contribute to the quantitative differences observed in clubroot resistance among Arabidopsis accessions.

**Fig. 5.**
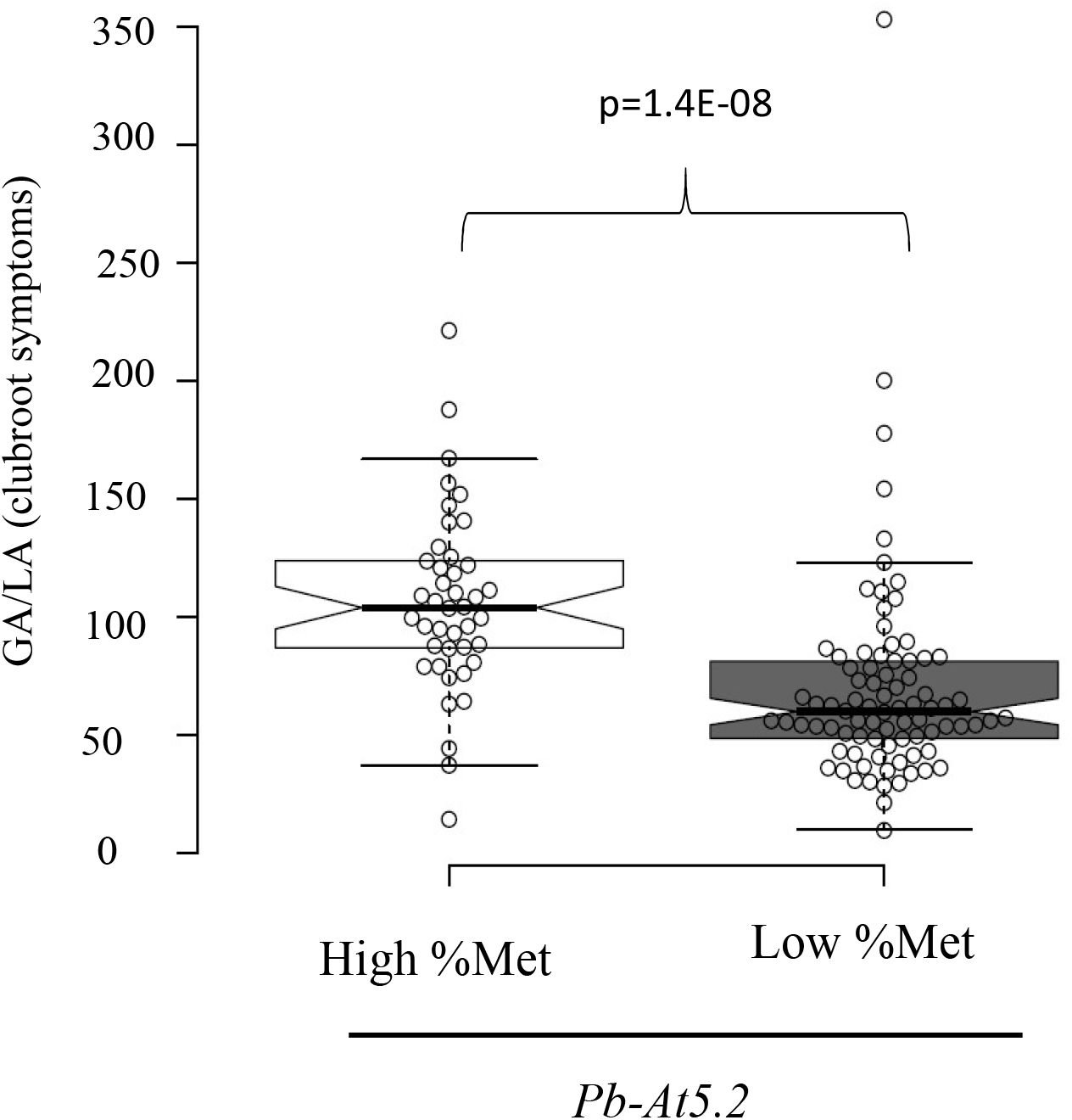
Clubroot symptom variation among Arabidopsis accessions is linked to epivariation at *Pb-At5.2*. Effect of *Pb-At5.2* epiallele variation on clubroot susceptibility, evaluated in a series of 126 Arabidopsis accessions harbouring similar Bur/Col-like genomic structure at the locus. In total 42 accessions have Col-0 like epiallele and 84 have Bur-0 like epiallele. For each accession, the mean GA/LA was obtained by modelling raw data of two biological replicates with two blocks (6 individual plants in each block). Center lines show the medians; box limits indicate the 25th and 75th percentiles as determined by R software; whiskers extend 1.5 times the interquartile range from the 25th and 75th percentiles, outliers are represented by dots; data points are plotted as open circles. The notches are defined as +/−1.58*IQR/sqrt(n) and represent the 95% confidence interval for each median. *p*-value (Wilcoxon test) is indicated.

### Pb-At5.2 epivariation is independent of cis- genetic variations

At the locus *Pb-At5.2*, the transposon AT5TE69050 was present in both parental genotypes, with no sequence variation that might have been the primary cause of the variation of DNA methylation on the two adjacent genes. Analysis of 34 out of the 287 accessions with the Col-0/Bur-0-like haplotype, did not reveal the presence/absence of TE insertion variant within the 26kb *Pb-At5.2* region (Quadrana et al., 2016; Stuart et al.), with the exception of a private helitron insertion in the accession NFA-10. Moreover, 18 and 16 of these accessions displayed hypermethylated and hypomethylated epialleles, respectively, indicating that variation in DNA methylation is not associated with TE presence/absence variants. In addition, the cis-nucleotide polymorphism located within the coding sequence of At5g47260 and detected in Bur-0 was absent in at least five others accessions sharing the hypomethylated epiallele (**Supplementary Figure S8**), indicating that the hypomethylated state of *Pb-At5.2* is not correlated with any specific DNA sequence polymorphism at the locus.

### The hypermethylated epigenetic variant is maintained by the RNA-independent pathway

Analysis of sRNAs identified in Col-0 (Stroud et al., 2014) revealed that the At547260/At5g47280 region is targeted mostly by 24-nt sRNA, which prompted us to generate sRNA profiles from non-inoculated roots of Col-0 and Bur-0 as well as to 7 and 14 dpi with *P. brassicae* isolate eH. Consistent with the pattern of DNA methylation observed before, we found high levels of sRNAs only in Col-0 (**Fig. 6a**). To explore further the mechanisms involved in the maintenance of methylation at this locus, we made use of publicly available methylomes of Col-0 mutant plants defective in one or several DNA methylation pathways (Stroud et al., 2013). Despite the high levels of sRNAs detected over the At547260/At5g47280 region, mutations affecting the RdDM pathway did not influence its DNA methylation level (**Supplementary Figure S9**). Conversely, DNA methylation was largely lost in mutants defective in sRNA-independent maintenance of DNA methylation, i.e. *ddm1*, *cmt2/3*, *met1*, and *suvh456.* (**Fig. 6b)**. These results raise the question of the role of the sRNA targeted to the Col-like hypermethylated region, whereas the methylation maintenance solely depends on the RNA-independent pathway.

**Fig. 6.**
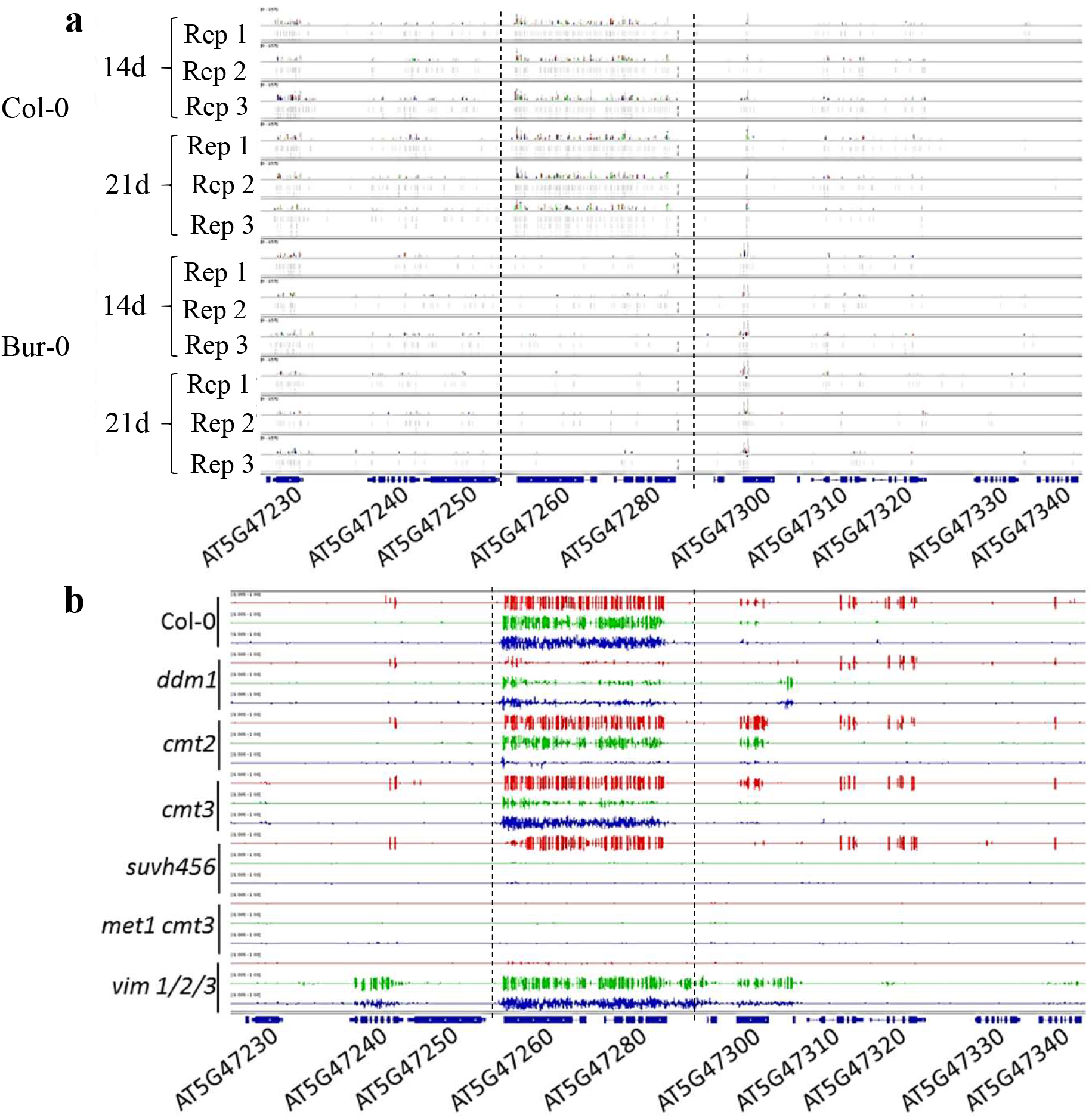
Epigenetic variation at *Pb-At5.2* correlates with the abundance of locus-targeted sRNA, but is maintained by the RNA-independent methylation pathway. **a** Mapping of sRNA-seq reads. Reads were obtained from roots of Col-0 and Bur-0 accessions at two time points at 14 and 21 days after sowing. For each condition, 3 bulks (numbered from Rep 1 to Rep 3) of 6 plants were used **b** Methylation state at the *Pb-At5.2* locus in knock-out lines (Col-0 genomic background) defective for the RdDM or non-RdDM pathways (Stroud et al., 2014). In red the methylation in CG context. In green the methylation in CHG context; in blue the methylation in CHH context.

## Discussion

To date, only a very small number of resistance QTL have been characterized at the molecular level (Nelson et al., 2017). Detection and fine mapping of resistance QTL is typically challenging not only because of the difficulties associated with measuring small variations in partial resistance in a large number of individual progeny, but also because resistance QTL can be sensitive to environmental changes (Laperche et al., 2017; Aoun et al., 2017; Aigu et al., 2018). However, technical issues may have only been part of the problem. Recent developments in the field of epigenetics suggest that the detection of some inherited resistance factors could be missed due to classic genetic approaches being based only on DNA sequence variation. In the present work, genome-wide association study failed at identifying any nucleotide variation in the 26 kb interval of *Pb-At5.2* associated with clubroot response. By contrast, clubroot resistance was clearly related to the epigenetic variation at two NLR-coding genes in this interval. This work thus uncovers for the first time an epigenetically driven expression polymorphism substantially contributing to the natural diversity of plant immune response.

Many examples of epialleles are metastable, *i.e.* they can be reversed by stochastic or non-identified factors (Weigel and Colot, 2012). Stability over multiple generations is a primary concern from both evolutionary and breeding perspectives. The epiallele described here seems to be extremely stable, as it was robustly detected in all our previous QTL investigations in *Arabidopsis*. This included two independent segregating progenies derived from Bur-0 and Col-0 (Jubault et al., 2008a) and additional studies with HIF lines 10499/13499 (Lemarié et al., 2015; Gravot et al., 2016). The high level of methylation and absence of At5g47260 and At5g47280 expression in Col-0 was found in a series of publicly available data obtained in different laboratories, with a diversity of plant tissues and conditions (Winter et al., 2007; Stroud et al., 2013; Klepikova et al., 2016; 1001 Genomes Consortium, 2016; Kawakatsu et al., 2016). It was also confirmed by our own data generated on inoculated roots, non-inoculated roots and leaf samples. Finally, this methylation pattern was also robustly found in multiple replicates of individual plants. Thus, *Pb-At5.2* can be unreservedly classified as a stable epialleles.

There has only been one recent report of a plant disease resistance caused by an inherited methylation variant affecting the expression of a resistance-related gene (Nishimura et al., 2017). In that study, a stable expression polymorphism (between Ler-0 and Ag-0 accessions) on the TIR-only encoding gene *RBA1* (At1g47370) affected ETI-responses to the *Pseudomonas syringae* effector *hopBA1*. This expression polymorphism was linked to the nearby presence/absence of a TE sequence in the promoter region of the gene and to *MET1-*dependent DNA methylation variation. However, because DNA methylation was reversed when the TE sequence was segregated away (**Figure S2** in Nishimura et al. (2017), DNA methylation variation is not epigenetic as it is an obligate consequence of sequence variation (i.e. presence/absence of the TE sequence). In the present study, we showed that DNA methylation variation in the region between At5g47260 and At5g47280, including the TE sequence *At5TE69050*, is not linked to any nucleotide/structural variation at the locus or elsewhere in the genome. Thus, *Pb-At5.2_COL_* and *Pb-At5.2*_BUR_ can be considered as ‘pure epialleles’, as defined by Richards (2006).

From available genomic and epigenomic data from the ‘1001 genomes project’, it can be extrapolated that the Bur-like clubroot resistance epiallele is present in about half of the accessions from the ‘Relict’, ‘Spain’, and ‘Italy/Balkans/Caucasus’ groups and 39 % of the accessions from the ‘North Sweden’ group (**Supplementary Data 3**). In contrast, the Bur-like epiallele is likely (taking into account missing methylation data) present at about 10 % in the ‘Germany’ group. On the other hand, the clubroot susceptibility of the Col-like epiallele was absent from the accessions in the ‘Relict’ and ‘North Sweden’ groups, but reached at least 16.8 % in the ‘Central Europe’ group. This sharp geographical structuration suggests that both epialleles confer fitness gains depending on environmental contexts. However, it does not appear to be obviously related to the distribution of clubroot incidence in *Brassica* cultures (usually low in the warm southern European regions). Keeping in mind that NLR can detect unrelated effectors from distinct microbial species (Narusaka et al., 2009), and echoing previous work (Karasov et al., 2014), we hypothesize that the maintenance of this epivariation in natural populations may reflect additional roles that *Pb-At5.2* plays against other plant pathogens (besides the control of clubroot infection).

At5g47260 and At5g47280 belong to a small heterogeneous cluster of three non-TIR-NLR located on chromosome 5, not far from the largest NLR-hot spot in the *Arabidopsis* genome (Meyers et al., 2003). None of the three genes have been previously shown to be involved in plant pathogen interactions. There are a few examples of tandem NLR genes coding for pairs of proteins that function as heterodimers (Cesari et al., 2014; Williams et al., 2014; Saucet et al., 2015). Similarly, the proteins encoded by these two epigenetically joint-regulated genes may function together in the control of cell defense responses during clubroot infection. Although the underlying molecular mechanisms are not known, the canonical example of the TIR-NLR heterodimer *RRS1*/*RPS4* corresponds to a recessive resistance locus, similar to *Pb-At5.2*. Among the 287 genotypes with Col/Bur-like genomic structure and available methylation data, both were genes were always methylated and it was thus not possible to use these genetic resources to gain information on the impact of silencing only one of the two genes. Additional molecular work (including CRISPR-Cas9-based nucleotide deletions or demethylation, in the Bur-0 accession, and the analysis of protein-protein interactions) is thus necessary to assess the individual role and possible partnership of these two genes.

Using public data (Zilberman et al., 2007), we found that 32 and 77 genes from the NLR family are methylated and unmethylated in Col-0, respectively (**Supplementary Data 5**). This suggests that plant genomes contain series of functional resistance genes, whose possible roles in biotic interactions are locked by epigenetic processes. This hypothesis is also supported by our recent study, where we demonstrated that the *ddm1*-triggered hypomethylation at different genomic loci resulted in unlocking genetic factors ultimately exerting a significant control over the development of clubroot symptoms (Liégard et al., 2019). It would now be interesting to carry out a careful genome wide analysis of methylation profiles of all NLR-genes among *Arabidopsis* accessions, which would take into account the structural diversity of all these individual genes (supported by additional targeted resequencing of NLR-loci). The intraspecific diversity of the methylation patterns of NLR and RLK/RLP genes in plants, their heritability, and their consequences on plant biotic interactions, may also deserve further attention in future studies.

## Methods

### Plant materials and growth conditions

Heterogeneous Inbred Family (HIF) lines 10499 and 13499 and their parental accessions Col-0 (186AV) and Bur-0 (172AV) were obtained from the Institute Jean Pierre Bourgin (INRA Versailles, France). *Arabidopsis thaliana* accessions were all purchased from the Nottingham Stock Center. The panel of 126 accessions was selected according to their methylation level at the region of interest (Kawakatsu et al., 2016). All accessions and in-house generated recombinant lines used in this study are listed in **Supplementary Data 1** and **Supplementary Data 3**. Seed germination was synchronized by placing seeds on wet blotting paper in Petri dishes for two days at 4 °C. Seeds were sown individually in pots (four cm diameter) containing a sterilized mix composed of 2/3 compost and 1/3 vermiculite. Growth chamber controlled conditions of 16 h light (110μmol m^−2^s^−1^) at 20 °C and 8 h dark at 18 °C were used to grow plants. The 126 *Arabidopsis* accessions and HIF were challenged by *P. brassicae* in two biological replicates in a completely randomised block designs (with two blocks per replicate, each block consisting in 6 plants by genotype). The *Arabidopsis* accessions Col-0, Bur-0 and the HIF 10499, 13499, 1381-2 and 2313-15 used in RNA-seq and smRNA-seq approaches were assessed when infected with *P. brassicae* or in uninfected condition in three randomised blocks. Almost all clubroot tests were performed with the isolate eH of *P. brassicae* described by Fähling (2003) which belongs to the most virulent pathotype P1. The resistance spectrum of *Pb-At5.2* was also assessed with a series of additional isolates Pb137-522, Ms6, K92-16 and P1^+^. For every isolate used in this study, the pathotype was validated in every experiment using the differential host set according to Some et al. (1996), also including two genotypes of *B. oleracea* ssp*. acephala* C10 and CB151. One ml of resting spore suspension (10^7^ spores.ml^−1^) prepared according to Manzanares-Dauleux et al (2000) was used for pathogen inoculation 10 days (d) after germination (stage 1.04 (Boyes et al., 2001)). This inoculum was applied onto the crown of each seedling.

### Phenotyping

HIF and *Arabidopsis* accessions were phenotyped three weeks after inoculation (21 days post inoculation (dpi) for their susceptibility to *P. brassicae*. Plants were thoroughly rinsed with water and photographed. Infected roots were removed and frozen in liquid nitrogen. Clubroot symptoms were evaluated through image analysis using the GA/LA index calculated according to Gravot et al. (2011).

### Fine mapping of the loci responsible for clubroot resistance

Fine mapping of *Pb-At5.2* was performed starting from crosses between HIF lines 10499 and 13499, followed by successive rounds of genotyping and clubroot phenotyping in subsequent plant generations (full details are given in **Supplementary Text S1**).

### RNA isolation, mRNA Sequencing and differentially gene expression determination

Total RNA was extracted from frozen and lyophilized roots (collected 14 days after inoculation) using the mirVana™ miRNA Isolation Kit (Invitrogen) according to the manufacturer’s instructions. After extraction, the RNA were quantified using a NanoDrop ND-1000 technologies and their quality was controlled using the RNA 6000 assay kit total RNA (Agilent). Samples with a RIN greater or equal to seven were used for sequencing. cDNA sequencing library construction and the sequencing were performed by the NGS platform at the Marie Curie Institute of Paris. Each library was paired-end sequenced on an Illumina HiSeq 2500 technology. Reads were aligned to the TAIR10 genome annotation and assembly of Col-0 *Arabidopsis thaliana* genome concatenated with the *P. brassicae* genome using STAR software (Dobin et al., 2013) (Version 2.5.3.a). Alignment conditions were selected according to the *Arabidopsis* genome. A maximum of five multiple read alignments were accepted and for each alignment no more than three mismatches were allowed. The resulting BAM files were used to determine read counts using the function Counts of Feature Counts software (Version 1.4.6) and the TAIR10 gff of *Arabidopsis* concatenated with the gff of *P. brassicae*. Differentially expressed genes were determined using the package EdgeR (Robinson et al., 2010) in R software (Team, 2013) (version 3.3.0). Raw counts obtained as described previously were used as entry data in EdgeR. Once CPM (Counts Per Million) was determined, only genes with at least one CPM in three samples were retained. Expression signals were normalized using the TMM method (Trimmed Mean of M-values) of the CalcNormFactors function of Edge R. Finally, the differentially expressed genes were determined using the decideTests function of Edge R using one minimum fold change between −1.5 and 1.5.

### GWAS analyses

A conventional GWA study on GA/LA data from 126 accessions was performed with easyGWAS (Grimm et al., 2017) (https://easygwas.ethz.ch/). Association analysis was performed with EMMAX (Kang et al., 2010) using 1 806 554 SNP with a MAF >0.05, after correction for population structure by including the three first principal components in the additive model.

### Small RNA isolation, sequencing, clusterization and differential presence determination

Small RNA was extracted from frozen and lyophilized roots (collected 14 days after inoculation) using the mirVana™ miRNA Isolation Kit (Invitrogen) according to the manufacturer’s instructions. After extraction, the small RNA was quantified using a NanoDrop ND-1000 and quality controlled using the Small RNA assay kit (Agilent). Samples with a RIN greater or equal to seven were used for sequencing. Construction of cDNA sequencing libraries and the sequencing were performed by the NGS platform of Marie Curie Institute of Paris. For each sample, single-end (50 bp) sequencing was carried out using Illumina HiSeq 2500 technology. Reads were aligned to the TAIR10 genome annotation and assembly of Col-0 *Arabidopsis thaliana* genome concatenated with *P. brassicae* genome using STAR software (Dobin et al., 2013) (Version 2.5.3.a), counts and clustered using Shortstack software (Axtell, 2013). The presence of differentially expressed sRNA was determined using the EdgeR (Robinson et al., 2010) package in R software (Team, 2013) (version 3.3.0). Raw counts obtained as described previously were used as entry data in EdgeR. Once CPM (Counts Per Million) were determined, only genes with at least one CPM in three samples were retained. Expression signals were normalized using the TMM method (Trimmed Mean of M-values) of the CalcNormFactors function of Edge R. Finally, the differentially expressed sRNA were determined using the decideTests function of Edge R using one minimum fold change between −1.5 and 1.5.

### RNA isolation and RT-qPCR analysis

Total RNA was extracted from the lyophilised root of accessions and HIF 21 days post infection using the trizol extraction protocol. Samples with residual traces of DNA were treated with DNAse (Promega ref M6 10A). Before reverse transcription of RNA to cDNA by the SuperScript II (Invitrogen), the RNA quality was verified by agarose gel electrophoresis gel. RT-qPCR was carried out in a LightCycler® 480 thermocycler (Roche) on cDNA obtained as described above. Gene expression was normalized using as reference two *Arabidopsis* genes defined as stable during infection using RNA-seq data (At1g54610, At5g38470) following Pfaffl’s method (Pfaffl, 2001). Primer sets were designed for each gene and are listed in **Supplementary Table S1**.

### CHOP PCR and qPCR assays

Gene methylation profiles were investigated using the enzyme McrBC (M0272L, BioLabs®)(Zhang et al., 2014). Forty ng of DNA was incubated with 0.5 μL BSA (20 mg/mL), 0.5 μL GTP (20 mM), 5 μL NEB (10X) and 0.2 μL McrBC (10000 U/mL). For CHOP PCR and qPCR 2 ng of digested and undigested DNA was used. For the CHOP PCR the temperature conditions were adjusted according to primer design and 35 amplification cycles were used. To determine the methylation state of the targeted region, each sample was digested or not (control) with McrBC before amplification. For the CHOP qPCR the temperature conditions were adjusted according to primer design and 30 amplification cycles of were used. Methylation levels of the target region were calculated as the percentage of molecules lost through McrBC digestion according to Silveira et al (2013) with the formula: (1-(2-(Ct digested sample - Ct undigested sample)))*100. The percentage of DNA methylation for At5g13440 and At5g47400, was calculated in all CHOP qPCR as controls. Primer sets designed for each gene are listed in **Supplementary Table S1**.

### Published data

The DNAseq data, RNAseq data, variant sequences and bisulfite data of the natural accessions studied were obtained from previous studies (1001 Genomes Consortium, 2016; Kawakatsu et al., 2016) archived at the NCBI SRA number SRP056687 and the NCBI GEO reference: GSE43857, GSE80744. The bisulfite data and small RNA data of *Arabidopsis* mutants studied were obtained from previous study (Stroud et al., 2014) archived at the NCBI GEO reference: GSE39901.

### Statistical analysis

Data were statistically analyzed using the R program (Team, 2013).

## Data availability

The data supporting the findings of this study are available within the paper and its supplementary information files. All unique materials used are readily available from the authors.

## Acknowledgments

We acknowledge the Versailles Arabidopsis Stock Center for providing HIF lines, the biological resource center BrACySol for furnishing Brassica accessions, the Gentyane Platform for their contribution in genotyping the fine-mapping population, the Institute Marie Curie for sequencing mRNA and sRNA. Cyril Falentin is acknowledged for his help in the design of Kaspar markers. IGEPP colleagues are acknowledged for their technical support for clubroot phenotyping and sampling.

## Author contributions

BL, AG, MMD and MJ designed and conducted the experiments. CL and AG carried out the fine mapping. CL, JL, JB, YA, BL, AG, MJ performed the phenotyping and sampling. BL, AG, JB, JL and MJ carried out epigenetic and gene expression studies. BL, YA, LQ and VC conducted bioinformatics analyses. AG, BL, LQ, VC, MMD and MJ participated in drafting and revisions of the manuscript.

