## Supplementary figures for "Natural Epiallelic Variation is Associated with Quantitative Resistance to the Pathogen *Plasmodiophora Brassicae*"

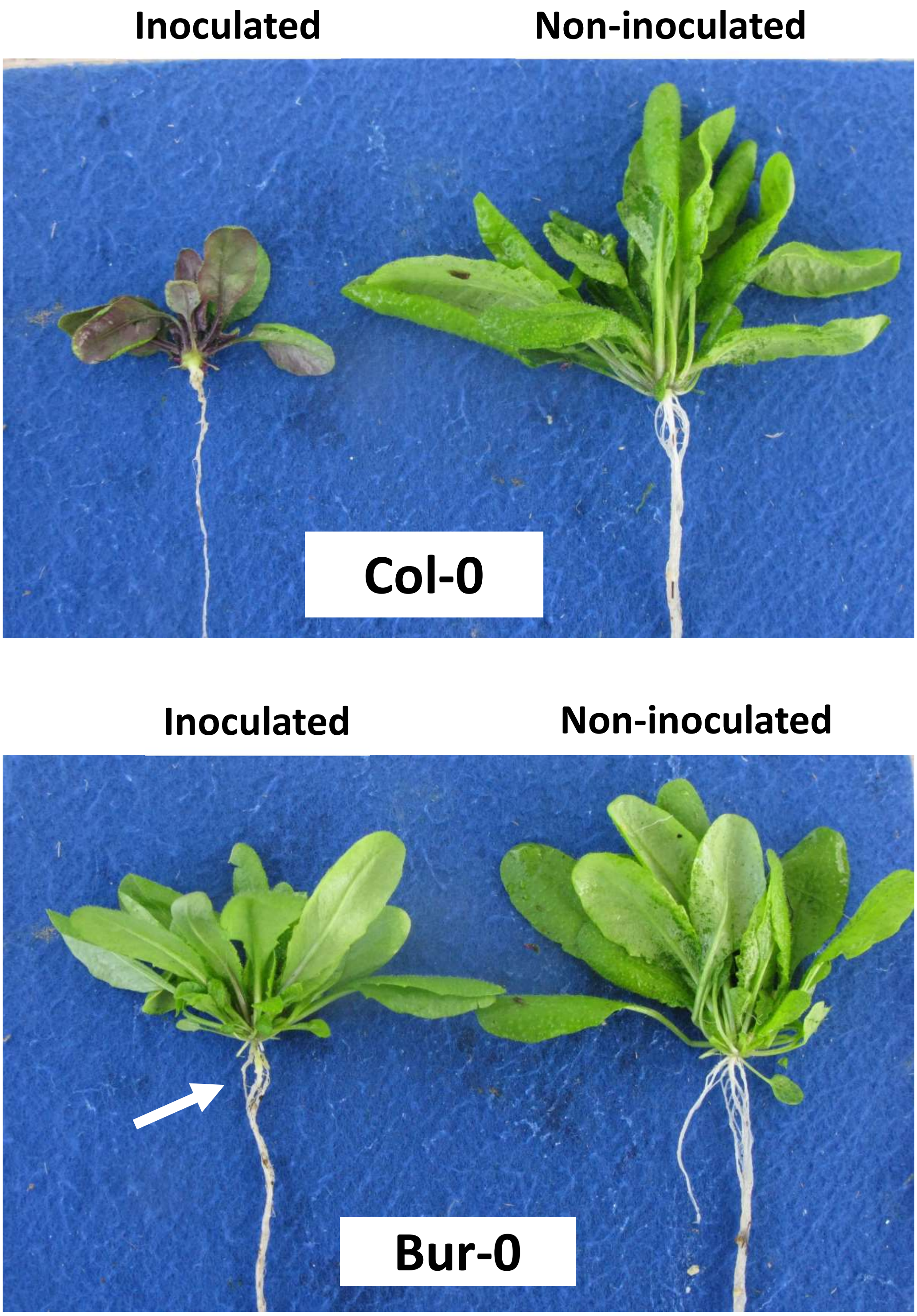


**Supplementary Figure S1 | Illustration of partial resistance to eH isolate in Bur-0, compared to the full susceptibility in Col-0.** Observations were done at 21 days post-inoculation. The white arrow indicates the presence of limited amount of galls in inoculated Bur-0, reflecting that partial resistance in this accession does not impair pathogen penetration but reduces its subsequent development. Plant individuals are representative of standard observations made in our experimental conditions. Levels of clubroot symptoms can vary depending on pathotests, but levels of clubroot symptoms are robustly found lower in Bur-0.

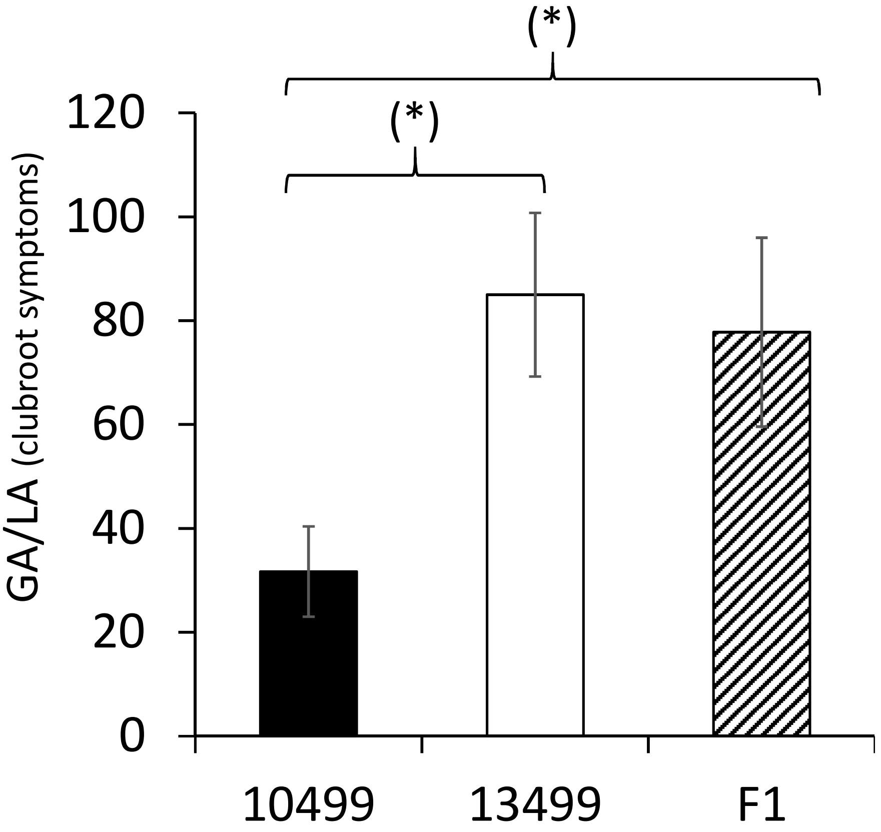


**Supplementary Figure S2 | Comparison of GA/LA disease index in the F1 progeny and in the parental lines 10499 and 13499.** GALA disease index is calculated through image analysis (details in the material and method part) from inoculated plants at 21 days post inoculation. Data are from 3 replicates (n=3). For each replicate, GALA disease index was calculated from 6 to 12 inoculated individual plants. Asteriks indicate statistical differences from the paired Student t-test (p=0.05)

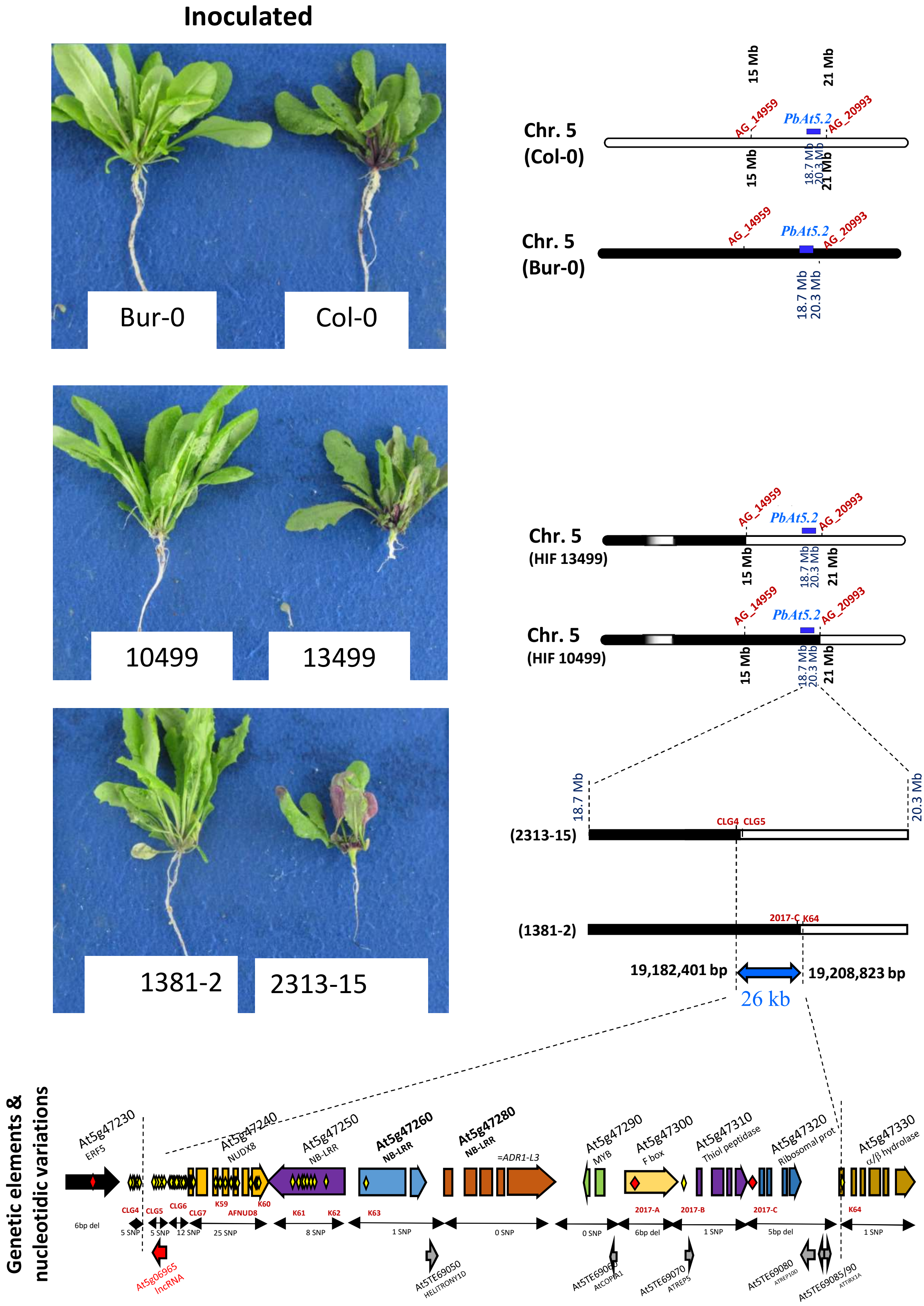


**Supplementary Figure S3 | Illustration of clubroot symptoms in a selection of genotypes used for the fine mapping.** Pictures of inoculated plants were taken at 21 days post inoculation. Genetic structure is indicated on the right side. Black=Bur allele, white=Col allele. Additional details in the legend of Figure 1.

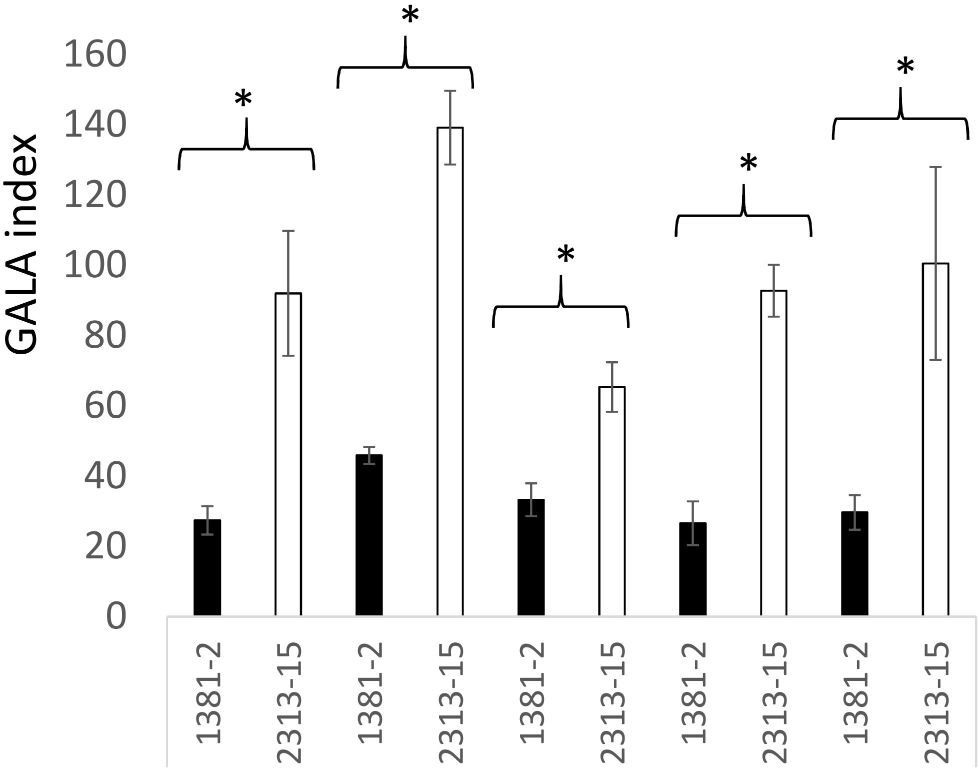


**Supplementary Figure S4 |** GALA disease index in RIL499-derived near-isogenic lines 1381-2 and 2313-15 challenged with a series of four European monospore *P. brassicae* isolates, and with the field isolate P1+. This last isolate is representative of emerging European pathotypes that are virulent on the variety ‘Mendel’ (a clubroot-resistant variety which has been cultivated at large scales in Europe and used as a source of resistance in many breeding programs). Data are means of 4 independent replicates (n=4). For each replicate, GALA values are means of 6 to 12 individual plants. Error bars indicate standard errors. Statistically different values (from Student T-test) are indicated by stars.

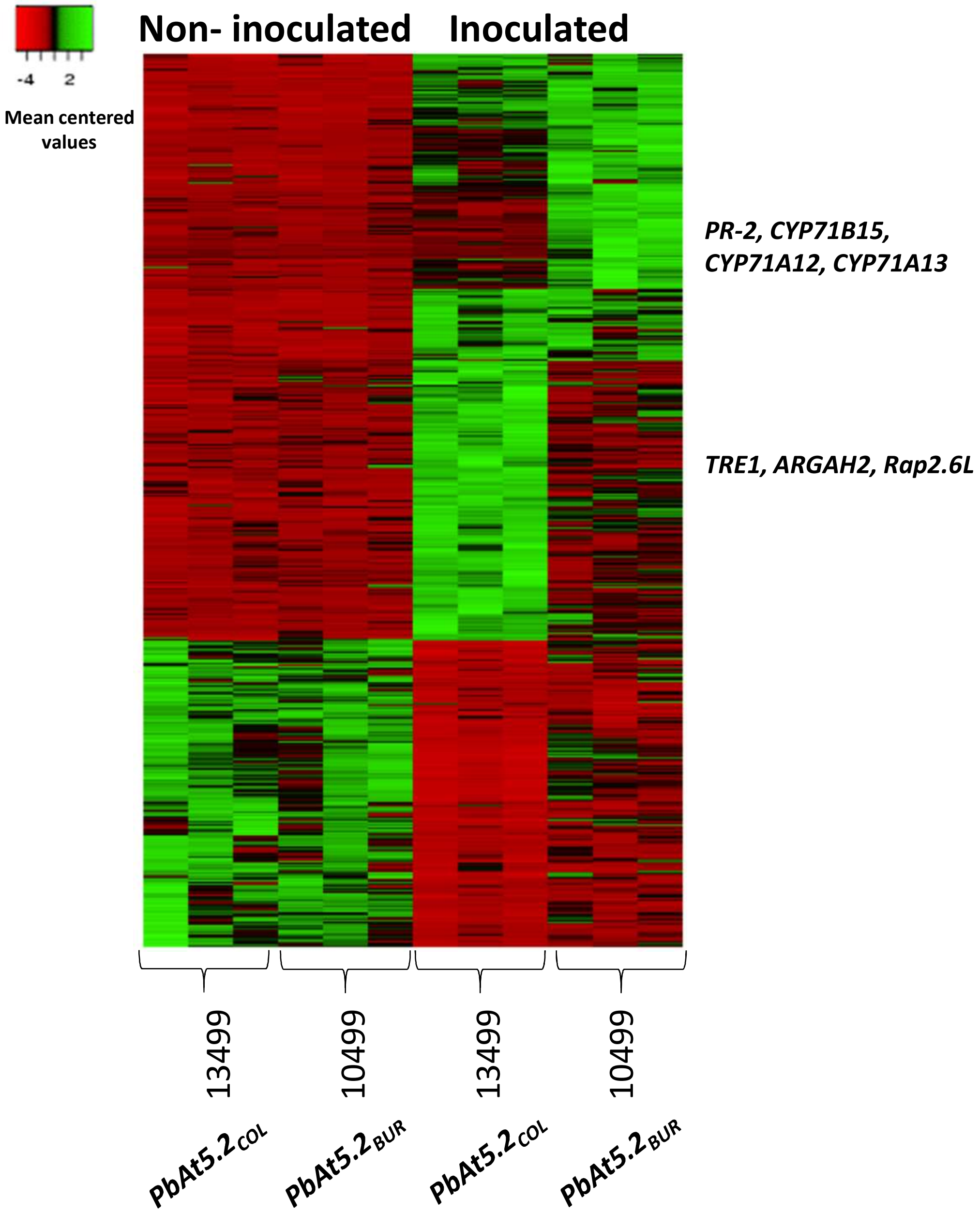


**Supplementary Figure S5 | Transcriptional regulations induced by isolate eH at 14 dpi in the recombinant HIF lines 10499 and 13499 harboring Bur or Col allele at *PbAt5.2*.** Data are mean-centered values of cpm. For each genotype, three columns are from three independent biological replicates. This set of 559 genes was selected as following: 1/Genes significantly induced (*p*-value<0.05 + log(FC)>2 or <-2) by eH isolate at 14 dpi in 10499 or 13499; 2/Mean gene expression>1. 61 genes, including *PR-2*, *CYP71B15*, *CYP71A12* and *CYP71A13*, were induced at higher levels in 10499 (*p*-value<0.05). 58 genes, including *TRE1*, *ARGAH2* and *Rap2.6L* were induced at higher levels in 13499 (*p*-value<0.05).

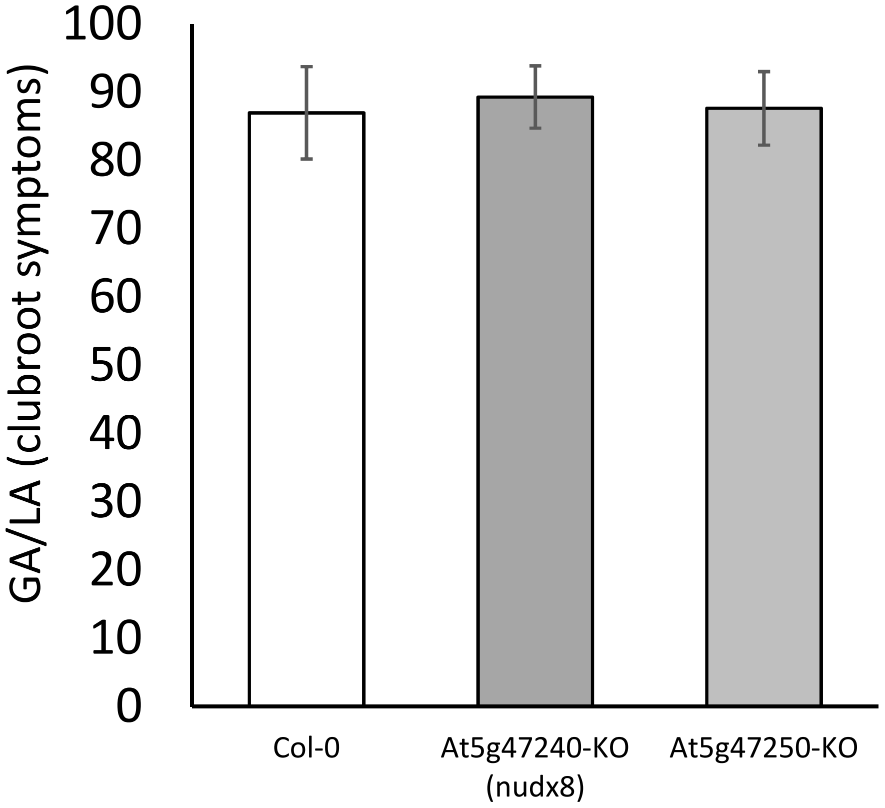


**Supplementary Figure S6 | GA/LA (clubroot symptoms) in the mutant lines defective for the expression of At5g47240 or At5g47250.** GA/LA was calculated through image analysis (details in the material and method part) from inoculated plants at 21 days post inoculation. Data are from 4 replicates (n=4). For each replicate, GA/LA was calculated from 7 to 11 inoculated individual plants. No statistical differences were found between the genotypes

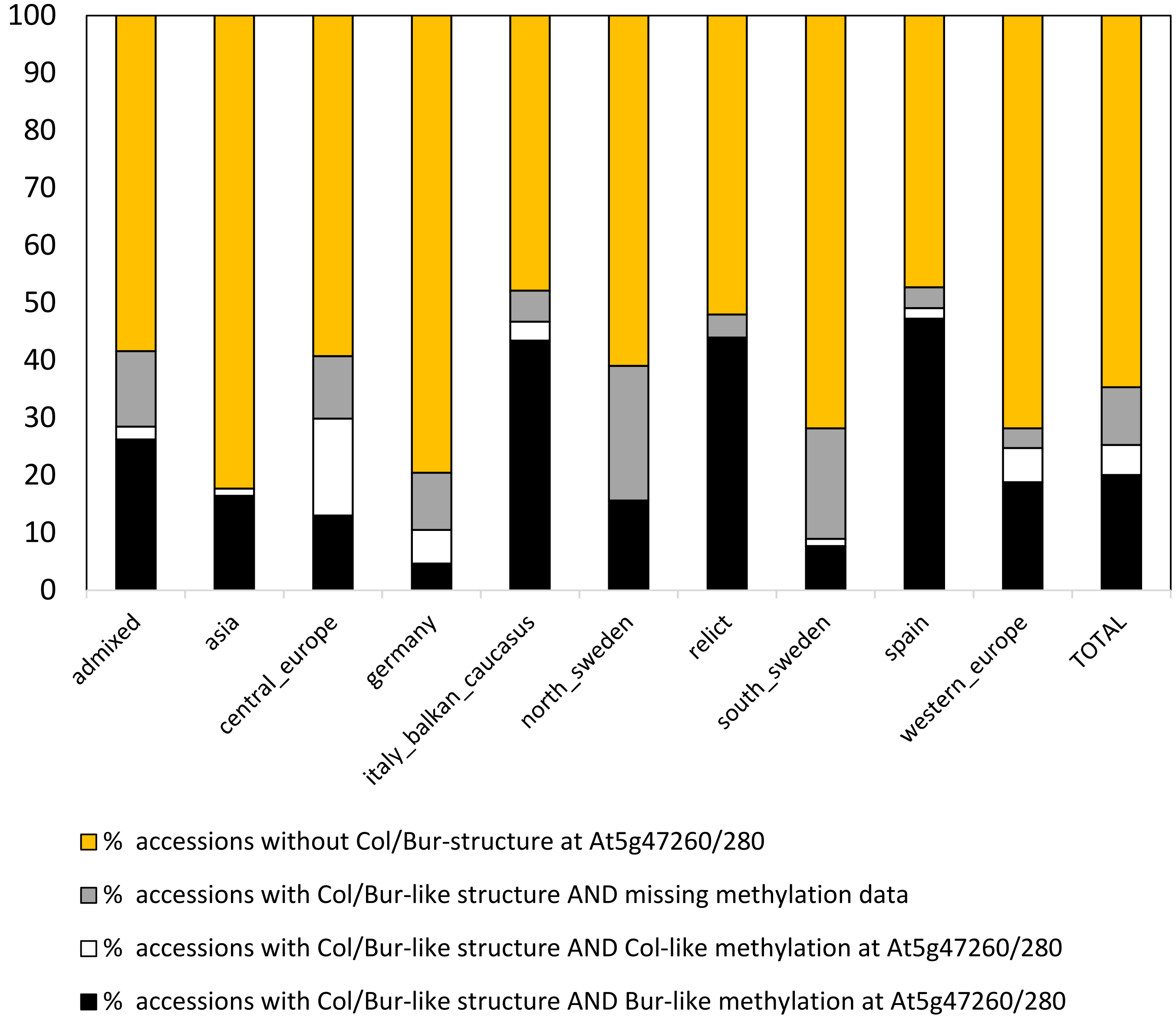


**Supplementary Figure S7 | Proportion of structural and epigenetic variations on the locus *PbAt5.2* among Arabidopsis accessions in each admixture group** (details in Supplementary Data 2).

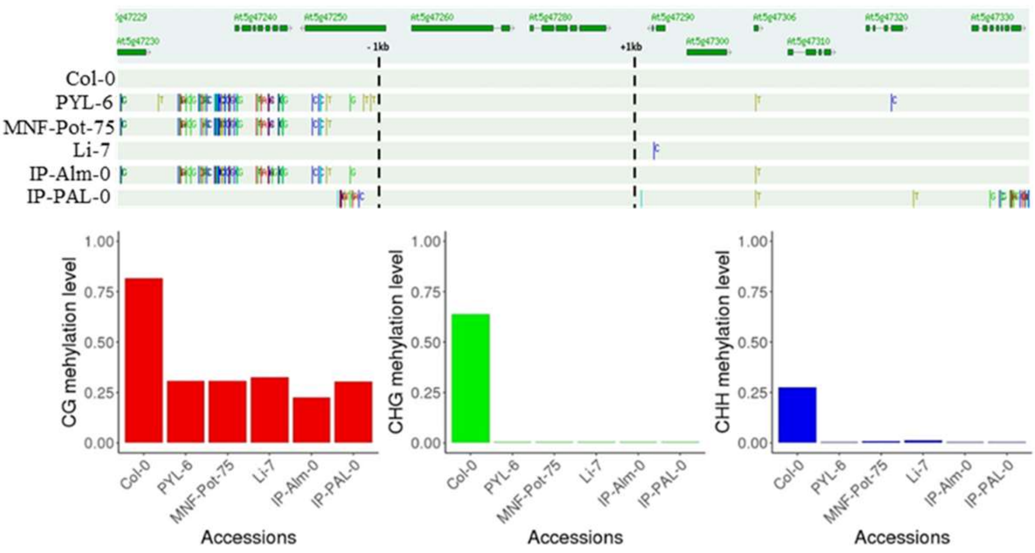


**Supplementary Figure S8 | The epigenetic variation on At5g47260 and At5g47280 is not related to SNP variations on At5g47260**. Sequence variant data were obtained from the signal SALK genome browser based on 1001 genome data. Average methylation level was calculated between 1 kb before the TSS site of At5g47260, up to 1 kb after the TSE site of At5g47280.

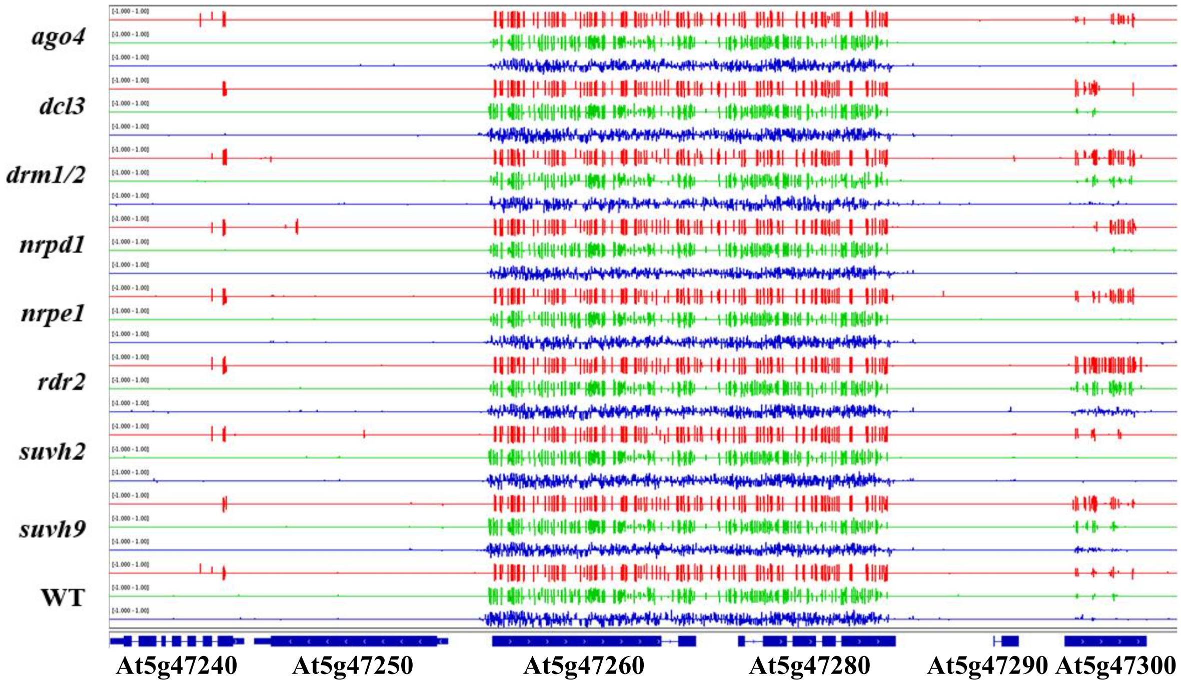


**Figure S9 | Methylation of *PbAt-5.2* region in Col-0 and in mutants in genes involved in RdDM methylation maintenance[51].** In red the methylation in CG context. In blue the methylation in CHG context; in green the methylation in CHH context. WT indicates the methylation profile of Col-0.
