## Supplementary information for "Natural Epiallelic Variation is Associated with Quantitative Resistance to the Pathogen *Plasmodiophora Brassicae*"

### Supplementary Text S1: Details of *PbAt5.2* fine mapping

#### Previous identification of *PbAt5.2*

A previous screen of *Arabidopsis thaliana* accessions found that Bur-0 is partially resistant to the *P. brassicae* eH isolate*,* whereas the canonical accession Col-0 is fully susceptible[1]. Two progenies were used to identify QTL controlling resistance to eH by linkage analysis: the first was derived from an initial Bur-0 (NASC accession N1028) x Col-0 (NASC accession N1092) cross followed by 6 generations of SSD (Recombinant Inbred Line set 20RV described in Simon et al. 2008). The second was derived from Col-0 x Bur-0 (described in[2]). These approaches led to the identification of a series of four additive QTL, including *PbAt5.2* ($R^2$=20%, Bur-0 allele at this locus confers partial resistance). The QTL peak of *PbAt5.2* was at the marker C5_19316 (around 19.3 Mb), with a confidence interval of 4.3 cM, between positions 18.7 Mb and 20.3 Mb (157 annotated genes between At5g46260 and At5g47690).

#### HIF499 for *PbAt5.2* validation

Among the RIL set 20RV from Simon et al. (2007), one RIL line (RIL499) displayed a single residual heterozygous region including markers c5_17570, c5_19313 and c5_20318, framed by the homozygous loci c5_14766 and c5_21319. Heterogeneous Inbred Family (HIF) lines 10499 and 13499 were derived from RIL499 to obtain homozygosity at this locus (Institut Jean Pierre Bourgin, INRA Versailles, France). These lines are near-isogenic, with identical combinations of Bur-0 and Col-0 homozygous genome sequences at every locus, except in the region between c5_14766 and c5_21319. The use of PCR-based markers (**Supplementary Data 1**) allowed us to reduce this interval between markers CL5_15283 and CL5_20983 (excluded). HIF line 10499 displayed a higher level of partial resistance to eH isolate compared to 13499, thus confirming the position of *PbAt5.2* in this interval ([3], see also the Figure 1 in the present work).

#### Generation and phenotyping of F1 individuals derived from crosses between 10499 and 13499

Fine mapping of *PbAt5.2* started from the cross between HIF-13499 (allele Col-0) and HIF-10499 (allele Bur-0) lines. Crosses were made in both directions, *i.e.* using one or the other parent as female. Heterozygosity in the *PbAt5.2* region was checked in several F1 individuals using a series of PCR-based markers (**Supplementary Data 1**). Clubroot index was evaluated in a series of F1 plants using the eH isolate, and was statistically identical to 13499, and higher than for 10499, thus suggesting that the Bur-0 resistant allele at *PbAt5.2* was recessive (**Supplementary Fig. 2**).

#### Screening of recombinant individuals in the segregating F2 progeny

One validated F1 plant was chosen from each of the two crosses. Those two plants were self-pollinized, and approximately 3200 F2 plants were sown (about 1600 from each cross). Individual F2 from number 1 to 1581 were from a 10499 x 13499 cross. Individual F2 plants from 1582 to 3153 were from a 13499 x 10499 cross. DNA was extracted from young leaves sampled from these 3152 plants, and then subjected to a first round of genotyping using a series of 10 KAPSPar SNP markers (list in **Supplementary Data 1**). Analyses were performed on the GENTYANE platform using a LightCycler 480 device (UMR INRA 1095, Clermont-Ferrand France). Due to low DNA concentrations in some samples, about 6% of the genotyping points were ‘negative’. Good or average-quality (i.e. genotyping ambiguity at maximum one marker) data were obtained for 2751 F2 individuals. Among those, 563 plants displayed at least one recombination in the region between At5g37660 and At5g51670, which represented about 20.5 % of the F2 plants, and was consistent with the distance of about 22 cM between those two marker genes. This mean value of 20.5 however masked a clear disparity between genotypes derived from the 10499 x 13499 cross (32% of individuals with one recombination in the region) from the 13499 x 10499 cross (18% of individuals with one recombination in the chromosomal region). Nevertheless, most of this disequilibrium was focused in the region between the markers on At5g42520 and At5g44630 (i.e. outside the confidence interval of the clubroot resistance *PbAt5.2* QTL), and the male x female direction of the initial hybridization step did not obviously affect the recombination rate in the region corresponding to the peak of the QTL (between At5g46910, At5g47120 and At5g47510).

**Phenotyping of recombinant F3 lines**

One hundred and seven recombinant F2 lines were selected based on the presence of a recombination event near the closest markers to the QTL peak (19.3 Mb). Using the seeds derived from the selfing of those individual lines, the clubroot disease index was estimated from 18 inoculated F3 plants (6 plants x 3 biological replicates). Those phenotyping data confirmed the presence of a segregating resistance locus in the region. Lines with Bur or Col homozygous alleles at markers At5g47120/At5g47510 displayed a mean GA/LA (clubroot symptoms) of 37.1 (SE=7.3) and 87.2 (SE=14.1), respectively. Lines with heterozygosity at those two markers displayed a mean disease index of 76.7 (SE=16), which was consistent with the above conclusion that the resistance Bur-0 allele was apparently recessive.

**High-density genotyping of recombinant F2/F3 lines**

A subset of 69 F2 recombinant lines was selected, based on the presence of a recombination event near the closest markers to the QTL peak (19.3 Mb). For each of these, 12 to 18 F3 progeny individuals were grown and their leaves were bulk-sampled, for subsequent analysis of 93 SNP (supplementary table 1). All leaf samples were analyzed at the GENTYANE platform. SNP genotyping was performed with the KASPar genotyping chemistry and Dynamic Array™ IFC 96*96 (UMR INRA 1095, Clermont-Ferrand France). Genotyping data obtained from bulked leaves of F3 individuals represents the genotypes of the parental F2 individuals. This genotyping workflow was also applied to a series of non-recombinant F2 lines and parental HIF 10499 and 13499 lines, which were used as controls. The comparison of the 93 SNP genotyping data and clubroot disease index for all those 69 lines finally led to the identification a small region between the markers K58=At5g47230prom (position 19,175,831 bp) and K65=AT5G47360 (position 19,214,446 bp). Due to the recessive status of the resistance allele, a series of recombinant lines with strategic recombination events (especially in the line 1381) were not exploitable.

**F4 lines with fixed alleles in the region of the resistance locus allowed further reduction of the *PbAt5.2* resistance locus interval**

DNA was extracted from 12 to 18 individual F3 plants derived from selfing a series of F2 recombinant lines. Among them, F3 lines with homozygosity in the region of the resistance locus were screened using the following PCR-based markers: CL5_16921=At5g42320; CL5_17802=At5g44200; CL5_18135=At5g44900; AF-NUD8; CL5_19601=At5g48375 (details in **Supplementary Data 1**). Homozygous F4 seed stocks were then obtained from the selfing of the selected homozygous F3 lines, and thereafter used for additional clubroot phenotyping assays. From the resulting data the interval could be reduced to a region between markers K58=At5g47230prom (position 19,175,831 bp) and K64 (position 19,208,823 bp), as shown in the **Figure 1E** (detailed phenotyping data also in **Supplementary Data 1**). Finally, every other SNP and indel in the region was analyzed by sequencing the PCR-amplified fragments from 2313-15, 1381-2, 2509-11, and 1600-5 (details of primers are given in the **Supplementary Data 1**). This allowed a final confidence interval of 26 kb between the marker CLG4 (19,182,401, in the promoter region of At5g47240), and the marker K64 (on SNP at position 19,208,823 bp, in At5g47330) to be identified.

#### Supplementary Text S2: Influence of *PbAt5.2* on transcriptomic responses to clubroot infection

The analysis of transcriptome responses to isolate eH, in 10499 and 13499, highlighted a series of 61 genes that were induced by infection only (or with higher range) in the presence of the resistance allele *PbAt5.2*$_{BUR}$ (**Supplementary Figure S6**). This series was enriched in genes associated with innate immunity, systemic acquired resistance, and notably included the SA-responsive gene *PR2*, and the set of genes *CYP71B15*, *CYP71A12*/*CYP71A13* involved in camalexin biosynthesis, confirming our previous studies on the cellular functions involved in clubroot resistance QTL *PbAt5.2*[4,5]. In contrast, a series of 58 genes was found to be induced by infection specifically (or with higher range) in the presence of the susceptibility allele *PbAt5.2*$_{COL}$. This set of genes included *ARGAH2*, a JA-regulated arginase encoding a protein involved in the biosynthesis of N-delta-acetylornithine, previously shown to play a role in basal resistance toward the eH isolate in genotypes harbouring the susceptible allele Col on QTL *PbAt5.2* [5,6]. This list also included the trehalase encoding gene *TRE1*, involved in resistance to massive amounts of trehalose synthesized by *P. brassicae* during clubroot infection[7,8].

**Table S1: List of primers used for qPCR and CHOP qPCR**

| Gene | Experimentation | LP primer (5' > 3') | RP primer (5' > 3') |
|---|---|---|---|
| At1g47550 | qPCR | CTCGCTCTTTCCGTCAAATC | CCCCAGTGTGAAAGTGCATC |
| At1g54610 | qPCR | GGTCGGACAGAGGTAGAGCAG | GTATGGTTCACGGGGTTTGT |
| At5g38470 | qPCR | TGTACTCGGGTATCCCTGCT | CTGGAGCTGCTGCTTGTTG |
| At5g47260 | qPCR | AAGGTGGTCCAATCGGGAAC | GATGGGGCAATCTGGTGTGA |
| At5g47280 | qPCR | AGTCTCTGGCTTGAGAGGGT | ATGGTCGAAGGTAGTTCCGC |
| At5g47260 | CHOP qPCR | TGCGTCGACCTATCGTTACA | CCATGCCGTATCAAGCAAC |
| At5g13440 | CHOP qPCR | ACAAGCCAATTTTTGCTGAGC | ACAACAGTCCGAGTGTCATGGT |
| At5g47400 | CHOP qPCR | GAAGCCGAACTGCAAACTGT | ATGGTCCGGCTCTAGGAAAA |
